# Apolipoprotein E genotype modulates longitudinal atrophy at the temporal lobe after mild traumatic brain injury

**DOI:** 10.1101/2022.08.13.503827

**Authors:** Shuoqiu Gan, Shan Wang, Yuling Liu, Yingxiang Sun, Kejia Liu, Xiaoyan Jia, Xuan Li, Ming Zhang, Lijun Bai

## Abstract

**Backgrounds:** Moderate or severe traumatic brain injury (TBI) is one of the strongest environmental risks for late-life dementia, especially Alzheimer’s disease (AD). However, the interaction between genetic factors and environmental exposure to mild TBI triggering neurodegenerative processes is still unclear. We used longitudinal imaging to differentiate spatial patterns of progressive brain volume loss in individual patients with and without carrying apolipoprotein E (APOE) ε4 carriers.

**Methods:** For 59 patients with acute mild TBI (mTBI, age: 36.92 ± 11.91 years; 14 APOE ε4 carriers) and 48 matched healthy controls (HCs, age: 37.92 ± 11.72 years; 10 APOE ε4 carriers), longitudinal (6-12 months follow-up) brain structure was assessed using voxel-based morphometry on T1-weighted scans. Longitudinal volume changes in the temporal lobe were characterized by the Jacobian determinant (JD) metric, reflecting spatial warping between the baseline and follow-up scans. JD values were regionally calculated and correlated with neuropsychological measures.

**Results:** MTBI patients lost a mean of 0.27% (SD = 0.43) of GM and 0.27% (SD = 0.50) of WM volume in the temporal lobe over the 6-12 month follow-up. Patients with a high genetic risk for AD (APOE ε4 allele carries) were associated with severer atrophy in the left superior temporal gyrus and middle temporal gyrus regions than the controls (*P* < 0.05). Furthermore, atrophy in these regions of gray matter could predict the performance change ratio of the language fluency in mTBI APOE ε4 carriers (*β* = 0.570, *P* < 0.05). While, compared with HC APOE ε4 non-carriers, mTBI non-carriers exhibited volume loss in the medial temporal lobe.

**Conclusions:** This prospective study provided evidence that the APOE ε4 allele interacting with mTBI increased the risk of AD phenotype development with language dysfunction.

**Graphic Abstract:** 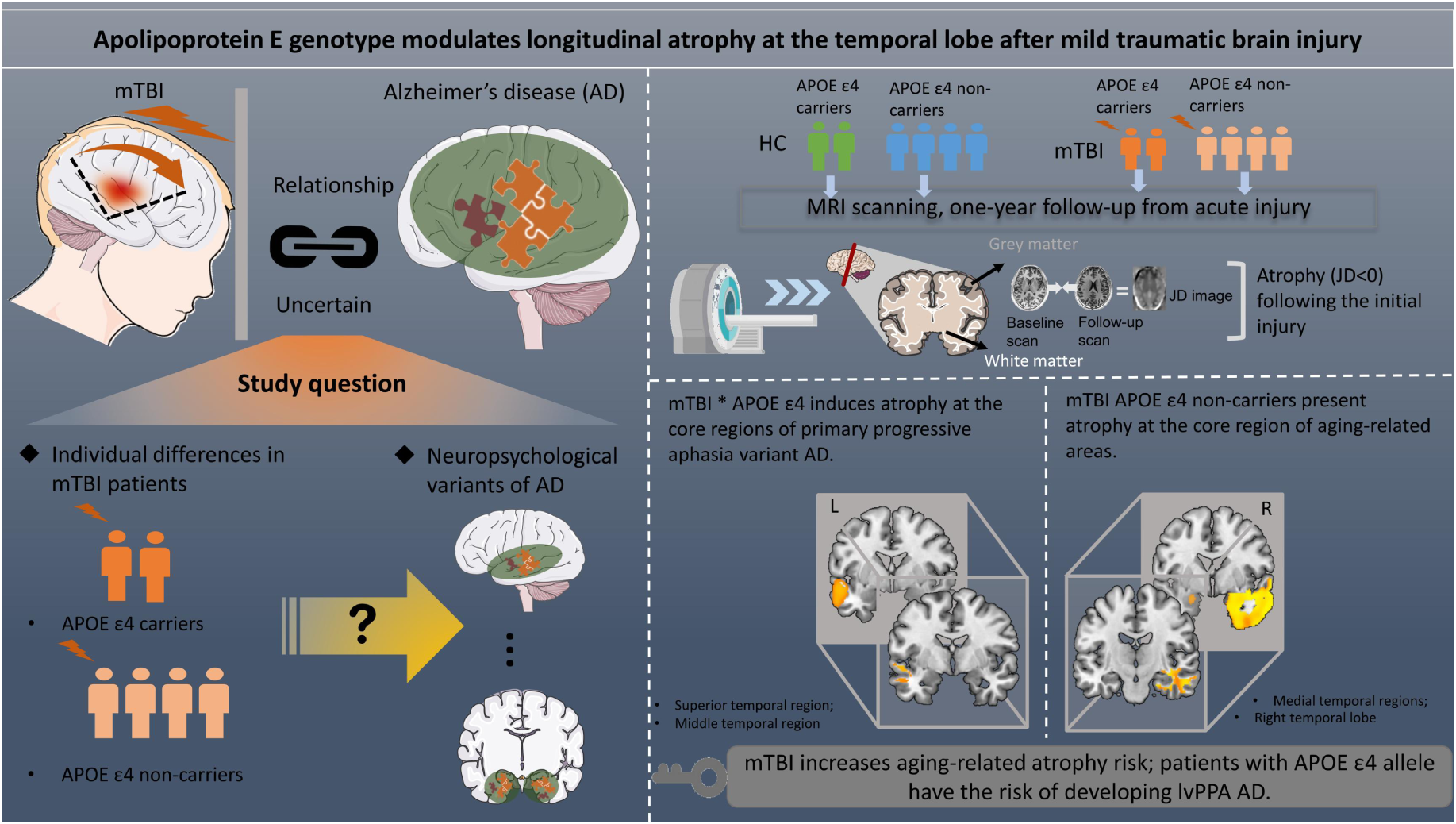

## Introduction

Traumatic brain injury (TBI) exposure has been identified as a risk factor for Alzheimer’s disease (AD) in later lifetime[1], even doubled the risk (odds ratio 2.29 [confidence interval 1.47–3.58])[2]. Indeed, TBI is ranked only after age, family history, and apolipoprotein E (APOE) genotype in importance as an AD risk factor[3], and the risk of AD from TBI is higher in APOE ε4 allele carriers than non-carriers [4-10]. TBI history interacting with APOE ε4 allele also leads to AD neuropathological changes, such as amyloid β-protein (Aβ) plaques deposition[11] and reduced cortical thickness in AD-vulnerable brain areas[12]. However, the interaction between genetic and environmental exposure of mild TBI triggering neurodegenerative process in a longitudinal way is still unclear.

Alternatively, some studies further suggest that TBI increases the risk for neurodegeneration only in certain sub-type[13, 14], and confer with AD phenotypes in specific domain such as language speech, executive function or memory[15, 16]. For instance, the atrophy of the temporal lobe is the most common neuropathological change shared by AD incident and TBI[17-19]. The initial atrophy in the lateral temporal lobe is also reported to be involved in the neurodegeneration pathway of several atypical AD variants which is symptomatic-related[20, 21], such as the language dysfunction[16]. However, such region-specific effects and increased risk of neurodegeneration in certain AD phenotypes needs further examinations.

To address this issue, we used a deformation-based morphometry method to directly model rates of brain atrophy in a voxel-wise frame [22, 23], and fits the repeated-measures nature of a longitudinal study with greater sensitivity and reducing biases from “asymmetric” imaging analyses[24, 25]. We hypothesized that the interaction of mild TBI (mTBI) exposure and high polygenic risk for Alzheimer’s disease would exhibit progressive atrophy in the identified AD-vulnerable regions. Such brain atrophy may present the region-specific effect and provide the evidence that TBI may increase the risk of neurodegeneration in certain AD phenotypes.

## Methods

### Participants

The flow of participants enrolled in the study was shown in the Figure 1. This study included 59 patients with acute mTBI (31 males, age: 36.92 ± 11.91 years) recruited from the local emergency department, and all the subjects had MRI scans at both acute phase (2.88 ± 2.43 day post-injury) and follow-up 6-12 months chronic phase (scan interval: 9.32 ± 3.76 months post-injury). The longitudinal MRI data (scan interval: 10.46 ± 3.20 months) was also acquired from 48 healthy controls (HCs, age: 37.92 ± 11.72 years; 22 males). Screening for the mTBI was based on the World Health Organization’s Collaborating Centre for Neurotrauma Task Force[26]. Inclusion and exclusion criteria were maintained for both samples and reported in our previous studies[27, 28]. HCs were enrolled through the advisements and carefully screened for any neurological or psychiatric disorders. All participants signed the written informed consent after the experimental procedures had been fully explained. The research procedures were approved by the local institutional review board and conducted in accordance with the Declaration of Helsinki.

**Figure 1.**
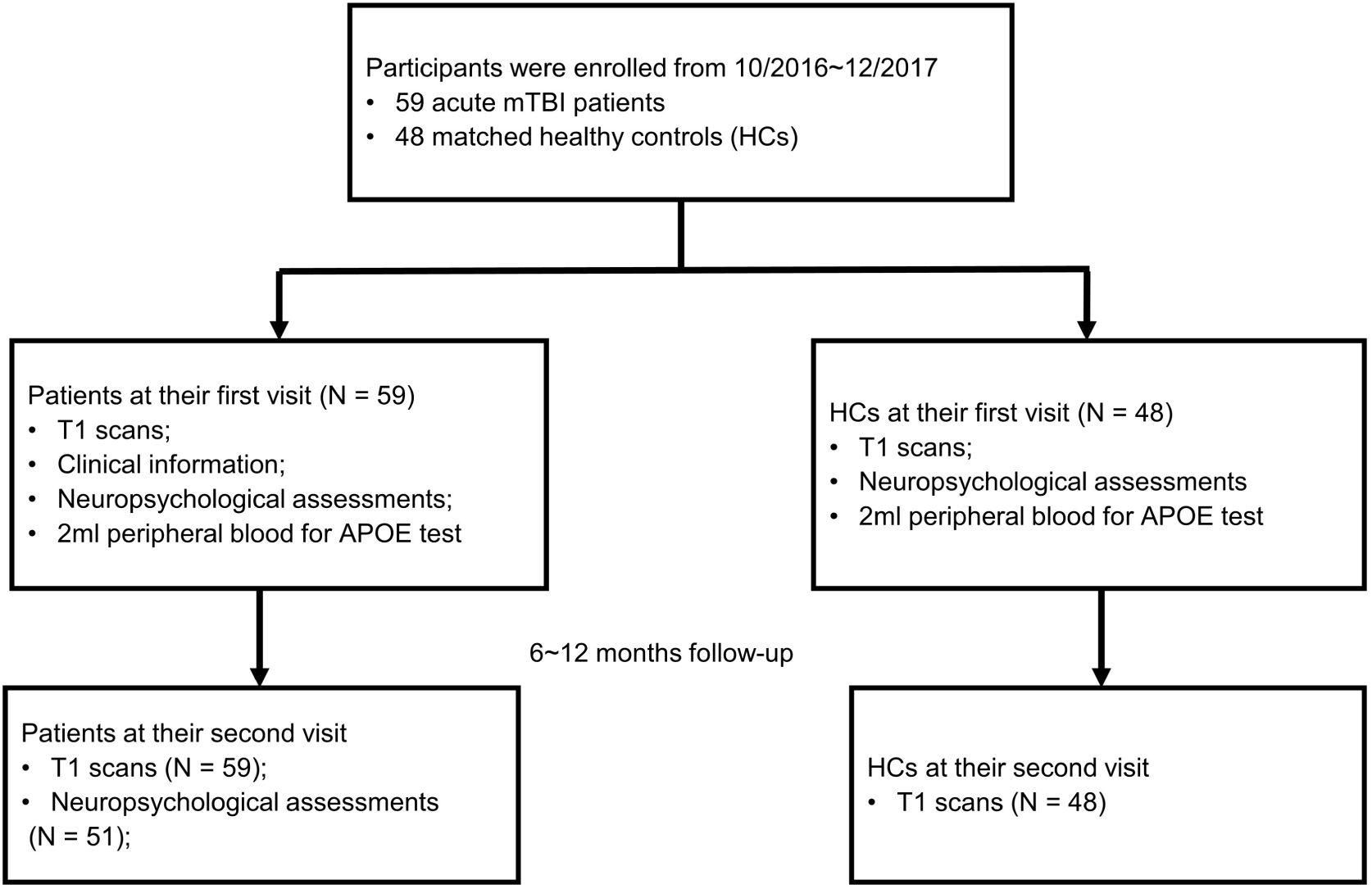
The flow diagram for the inclusion procedure of participants. Abbreviations: APOE = apolipoprotein E; mTBI = mild traumatic brain injury; HCs = healthy controls.

### Clinical symptom and neuropsychological assessments

Clinical assessments were performed within 48 hours of MR imaging for all the participants. The post-concussive symptoms (PCS) was evaluated by the International Classification of Diseases, 10th edition clinical (ICD-10) criteria[29]. Patients were classified as having PCS (PCS+) or not (PCS-) according to the split criteria of whether they had three or more symptoms in the PCS. The anterograde amnesia duration of patients was also recorded at their first visit.

Neuropsychological assessments were conducted at both the initial and follow-up visits for patients, while only at the first visit for healthy comparison. We focused on information processing speed (IPS) and language performance in this study, because they were the most prominently affected cognitive domain in AD[30, 31]. Information processing speed was evaluated by the Digit Symbol Coding test (DSC)[32] and Trail-Making Test (Part A, TMT-A)[33]. The language impairment was identified by using the forward digit span and language fluency test[16]. Z-score composed from these two tests in each domain was summarized as the performance of IPS and language, with lower scores indicating worse performances.

### APOE Genotyping

Blood samples were collected from all the subjects for APOE genotype analysis. DNA was extracted from peripheral blood samples using Omega D3494-01 Blood DNA Midi Kit (Omega Bio-tek, Inc., United States). Two single-nucleotide polymorphisms (SNPs; rs429358 and rs7412) were genotyped to identify APOE genotypes including the APOE-ε2, -ε3, and -ε4 alleles using Assay Design Suite v2.0 (Agena Bioscience, Inc., San Diego, CA). All genotypes containing the ε4 allele (ε4/ε4; ε4/ε3; ε4/ε2) were combined as the composite APOE ε4+, and others (ε2/ε2; ε2/ε3; ε3/ε3) were combined as the composite APOE ε4-. APOE genotype details were listed in the Table 1.

**Table 1.** Demographic, clinical and neuropsychological of patients and healthy controls.

| Characteristics | mTBI<br>(N = 59) | HC<br>(N = 48) | *P* value |
| --- | --- | --- | --- |
| Age (Years) | 36.92 ± 11.91 | 37.92 ± 11.72 | 0.664 |
| Sex (Male/female) | 31/28 | 22/26 | 0.490 |
| Education (Years) | 8.85 ± 3.91 | 10.47 ± 5.77 | 0.098 |
| APOE (+ε4/-ε4) | 14/45 | 10/38 | 0.721 |
| Scan interval (Years) | 0.78 ± 0.31 | 0.87 ± 0.27 | 0.099 |
| Injury causes |  |  |  |
| Traffic accident (%) | 55.93 | - | - |
| Fall (%) | 8.48 | - | - |
| Assault (%) | 30.51 | - | - |
| Other (%) | 5.08 | - | - |
| Injury severity |  |  |  |
| PCS (+/-) | 48/11 | - | - |
| Coma duration (Minutes) | 12.14 ± 11.07 | - | - |
| AA duration (Hours) | 0.17 ± 0.49 | - | - |
| IPS (Z-score) | -0.18 ± 0.84 | 0.22 ± 0.99 | 0.014* |
| DSCT | 36.33 ± 14.71 | 44.35 ± 17.63 | 0.006* |
| TMT_A (Seconds) | 59.09 ± 42.25 | 47.75 ± 29.45 | 0.059 |
| Language speech (Z-score) | -0.15 ± 0.84 | 0.19 ± 0.83 | 0.018* |
| Language fluency | 16.71 ± 5.24 | 18.79 ± 5.80 | 0.027* |
| Digital span (Forward) | 7.78 ± 1.53 | 8.25 ± 1.45 | 0.054 |

Note.— *P* values were derived from with the $\chi^2$ test, independent T-Test or Mann-Whitney U test, as appropriate. Abbreviation: HCs = healthy controls; mTBI = mild traumatic brain injury; APOE +ε4 = with APOE ε4 allele; APOE -ε4 = without APOE ε4 allele; PCS + = with post concussion symptom complaints; PCS - = without post concussion symptom complaints; AA duration = anterograde amnesia duration; IPS = information processing speed; DSCT = digit symbol coding test; TMT\_A = trail-making test part A
\* $P$ value indicates a significant difference at $\alpha = 0.05$ .

**Table 2.** Linear regression of neuropsychological performance changes with atrophy of APOE ε4-vulnerable ROIs.

| Variables | Model 1 |  | Model 2 |  | Model 3 |  | Model 4 |  |
| --- | --- | --- | --- | --- | --- | --- | --- | --- |
| | $\beta$ | $P$ | $\beta$ | $P$ | $\beta$ | $P$ | $\beta$ | $P$ |
| JD value of GM | 0.57 | 0.033* | 0.56 | 0.048* | 0.56 | 0.059 | 0.42 | 0.19 |
| JD value of WM | -0.24 | 0.417 | - | - | - | - | - | - |
| Sex |  |  | 0.072 | 0.78 | 0.075 | 0.78 | 0.29 | 0.42 |
| Age |  |  |  |  | -0.018 | 0.95 | -0.31 | 0.46 |
| Education |  |  |  |  |  |  | -0.44 | 0.36 |

Note.— JD value of GM = JD value in mTBI-APOE ε4 vulnerable ROI of grey matter; JD value of WM = JD value in mTBI-APOE ε4 vulnerable ROI of white matter
\* $P$ value indicates a significant difference at $\alpha = 0.05$ .

### Neuroimaging Data Acquisition and Preprocessing

High-resolution structural T1-weighted images for both acute and follow-up examinations were acquired using 3T GE 750 MRI with a 32-channel head coil. The high-resolution sagittal three-dimensional T1 image was collected using BRAVO sequence: repetition time = 8.15 ms, echo time = 3.17 ms, slice thickness = 1 mm, field of view = 256 × 256 mm, matrix size = 256 × 256, flip angle = 9°. Data processing procedure was showed in the Figure 2. Here, we used SPM12 longitudinal registration[25] to obtain individual JD image (Figure 2A-B). Following the pipeline in SPM12, the individual JD image and “temporal average” image in native space were generated by iteratively co-registering baseline and follow-up images. In this step, the time interval between two scans of each subject was weighted on their JD image to adjust the longitudinal changes with differing inter-scan intervals. JD value in each voxel representing the geometric warping amount between baseline and follow-up images. The positive and negative values represented an increase or decrease in size at the follow-up comparing with the baseline. Then, the individual “temporal average” image was segmented into grey matter (GM) and white matter (WM). The study-specific longitudinal template was defined in 40 randomly-selected samples (including 20 mTBI patients and 20 HCs) by using DARTEL (Diffeomorphic Anatomical Registration using Exponentiated Lie algebra) for registration use. Next, the GM and WM tissue segmentation images were multiplied by JD images to generate tissue-specific JD images. These images were registered to Montreal Neurological Institute (MNI)-152 space with resampling to 1.5mm^3^ voxels, via study-specific longitudinal template using DARTEL. Then smoothing with an 8mm full-width half maximum kernel was used and the data were modulated to preserve tissue amounts, by which the normalized GM and WM JD images were obtained for the following voxel-wise statistical analysis. Considering the temporal lobe being vulnerable for both mTBI and AD, the voxel-wise group comparisons of JD images were limited within the GM and WM of bilateral temporal lobe respectively, which were defined with the MNI Structural atlas (Figure 2C). In order to examine whether the longitudinal atrophy of temporal lobe was specifically modulated by APOE ε4 allele, we additionally employed the whole-brain voxel-wise group comparisons.

**Figure 2.**
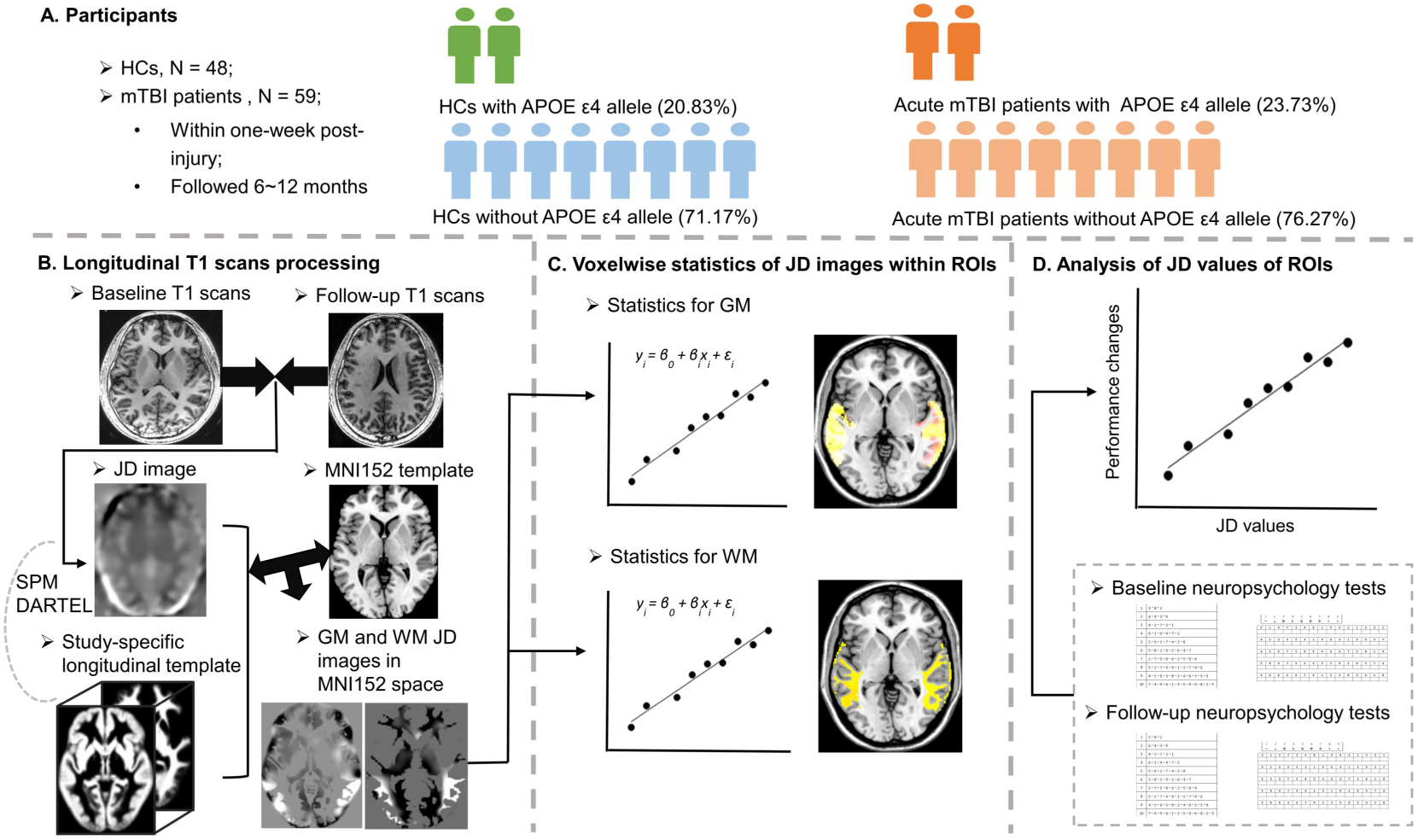
Overview of study design. (A) Participants recruited in the present study. Fifty-nine patients with acute mild TBI (mTBI, 14 APOE ε4 carriers) and 48 healthy controls (HCs, 10 APOE ε4 carriers) were recruited in the present study. All the participants were followed 6∼12 months. (B) Preprocessing of longitudinal analysis on T1-weighted image entailed an initial symmetric within-subject registration for each participant’s baseline and follow-up scans. This generated a within-subject ‘temporal average’ and Jacobian determinant (JD) image, representing the voxel-wise spatial expansion or contraction necessary to match baseline and follow-up images. Average images were then segmented into grey and white matter. A random selection of 20 mTBI patients and 20 HCs was used to define a study-specific longitudinal template with DARTEL for the following registration use. Individual average images and JD images were then normalized (smoothed by 8mm full-width half maximum kernel) to MNI152 space with resampling to 1.5mm^3^ voxels via the longitudinal template using DARTEL. (C) Longitudinal analysis included voxel-wise group comparisons using Randomise in FSL. In the current study, the voxel-wise group comparisons of JD images were limited in the bilateral temporal lobe extracted according to MNI Structural atlas. (D) Within the APOE subgroup of patients, multivariable linear regression analysis was conducted between the JD values regionally and the ratio of neuropsychological performance changes.

### Statistics

Voxel-wise statistical analysis was conducted using FSL Randomise software[34] for two-way between-subjects (group: HC vs. mTBI; APOE: APOE ε4+ vs. APOE ε4-) analysis of covariance in a generalized linear model (GLM), with the age, sex and education as covariates. Multiple testing correction used 5,000 permutations to generate statistically-corrected voxel-wise P-values for each contrast, with threshold-free cluster enhancement (TFCE) correction (*P* < 0.05). Genotype-vulnerable regions in patients was defined by each APOE subgroup (i.e., APOE ε4+ and APOE ε4-) comparison between HCs and mTBI, where mTBI-APOE ε4+/ε4-vulnerable ROIs in GM and WM were obtained.

Analysis of demographic, neuropsychological, clinical measures and relationships between ROI-based JD values with longitudinal changes of neuropsychological tests were performed in the SPSS (version 21.0; IBM, Armonk, NY). All the analyses for continuous variables were based on the normality distribution test. The group differences for demographics, APOE (ε4+ and ε4-), clinical and neuropsychological assessments were compared by using Chi-square analyses or Mann-Whitney U tests. Within each APOE subgroup of patients, to evaluate whether the longitudinal atrophy was symptomatic-related, the linear regression of neuropsychological performance change was performed with mean JD values of genotype-vulnerable regions.

## Results

### Demographic, clinical and neuropsychological assessments

Demographic, clinical and neuropsychological information for all participants were listed in the Table 1. Patients with mTBI and HCs were matched for age (*t* = 0.44, *P* = 0.66), sex (*χ*^2^ = 0.48, *P* = 0.49) and education level (*W* = 1679, *P* = 0.10). The proportion of mTBI cases carrying APOE-ε4 allele (23.73%) was not significantly different to the proportion of HCs (20.83%) (*χ*^2^ = 0.13, *P* = 0.72). Scan interval of mTBI patients (0.78 ± 0.31, years) was also matched with HCs (0.87 ± 0.27, years) (*t* = 1.65, *P* = 0.10). Mild TBI patients showed worse performance in the IPS (*t* = 2.23, *P* = 0.014) and language fluency (*t* = 2.12, *P* = 0.018) at their acute phase, compared with HCs at their first visit. Additionally, patients with APOE ε4 allele showed no significant differences to patients without ε4 allele in performance of IPS (*t* = 0.94, *P* = 0.35) and language fluency (*t* = 1.27, *P* = 0.21). Comparing with the initial neuropsychological assessments, mTBI patients obtained no significant recovery at follow-up in both IPS (*t* = 0.60, *P* = 0.551) or language fluency (*t* = 1.72, *P* = 0.089).

### Longitudinal atrophy of the temporal lobe following mTBI

Nearly 10 months post-injury follow-up, mTBI patients lost a mean of 0.27% (SD = 0.43) of the temporal lobe gray matter volume, and 0.27% (SD = 0.50) of the temporal lobe white matter volume. Healthy controls lost 0.07% (SD = 0.40) of the temporal gray matter volume, while gained 0.04% (SD = 0.46) of the temporal white matter volume over the same period.

Between-group comparison in Jacobian determinant image exhibited greater voxel-wise volume reductions of the temporal lobe during near one-year post-injury of mTBI, compared to controls (Figure 3). For the gray matter, a significant longitudinal atrophy was observed in the bilaterally temporal regions with saliently left-lateralization in mTBI patients, mainly containing extensive area of the left medial and lateral temporal lobe, compared to controls (Figure 3A; *P* < 0.05, TFCE corrected). For the white matter, mTBI patients showed a progressive atrophy mainly at the left lateral temporal regions and right middle temporal gyrus, compared to controls (Figure 3B; *P* < 0.05, TFCE corrected).

**Figure 3.**
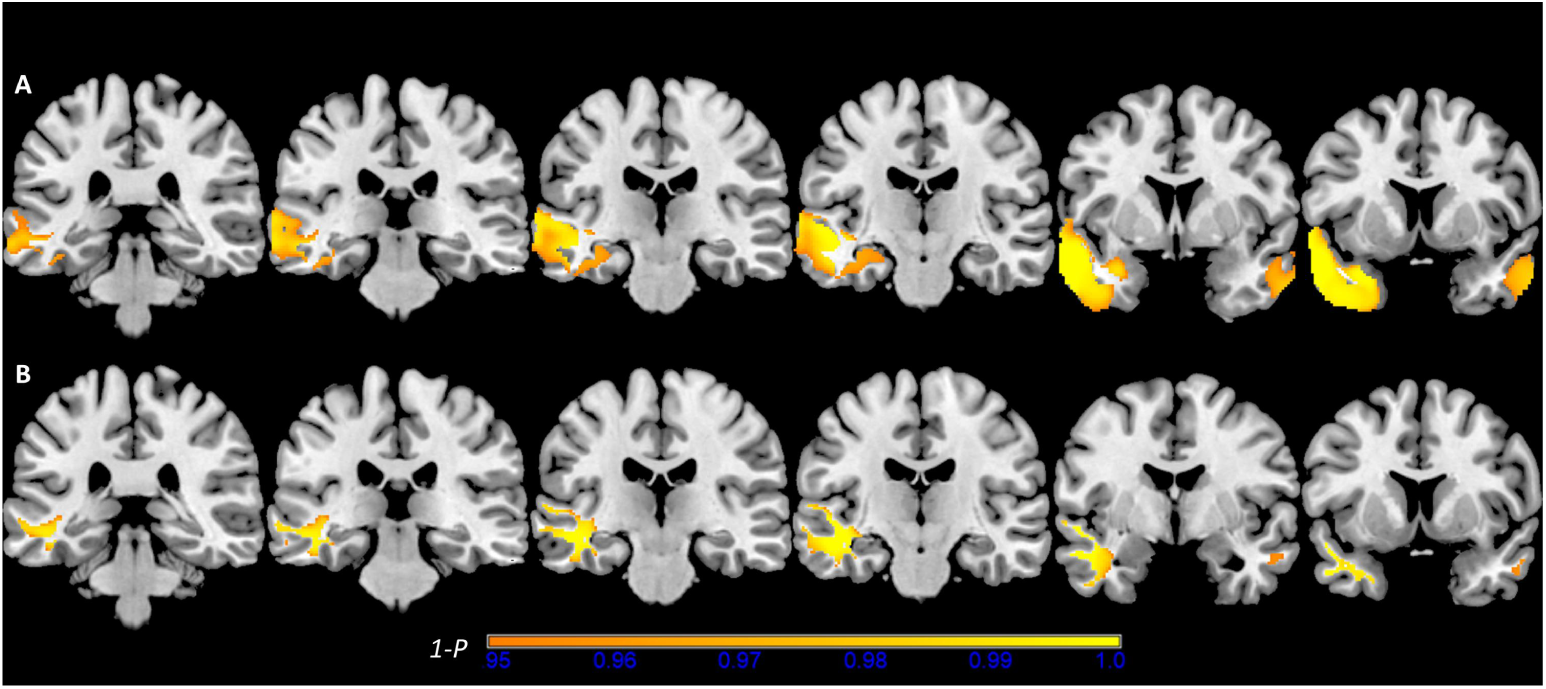
Patients with mTBI showed extensive longitudinal atrophy in the temporal lobe comparing with HCs. (A) Atrophy areas of mTBI patients in the grey matter of the temporal lobe. Patients exhibited the longitudinal grey matter volume loss distributing at the bilateral temporal lobes, with a significant left-lateralization atrophy pattern containing both the medial and lateral temporal grey matter area. (B) Atrophy areas of mTBI patients in the white matter the temporal lobe. Patients exhibited the longitudinal white matter volume loss mainly distributing at the left temporal lobe. Results are thresholded at voxel-wise *P* < 0.05, with threshold-free cluster enhancement (TFCE) correction.

### Longitudinal atrophy spatial pattern in mTBI with different APOE genotype

To find out the discriminative longitudinal atrophy pattern of APOE genotype individuals after mTBI, a voxel-wise between-group comparison of the temporal lobe JD images between the mTBI and the HC was also conducted for APOE ε4 carriers and non-carriers respectively.

The mTBI APOE ε4 carriers exhibited the marked atrophy in the left superior temporal gyrus and middle temporal gyrus for both gray and white matter (Figure 4A and 4B; *P* < 0.05, TFCE corrected), comparing with the healthy control ε4 carriers. While, mTBI patients without APOE ε4 allele showed a right-lateralization longitudinal atrophy pattern. Comparing with the ε4 non-carrier healthy control, the greater atrophy in patients without ε4 allele was primarily located in the gray matter of the right lateral temporal and bilateral medial temporal areas (i.e., hippocampus and parahippocampus) (Figure 4C; *P* < 0.05, TFCE corrected), and the longitudinal atrophy in the white matter of the right lateral and medial temporal cortex (Figure 4D; *P* < 0.05, TFCE corrected).

**Figure 4.**
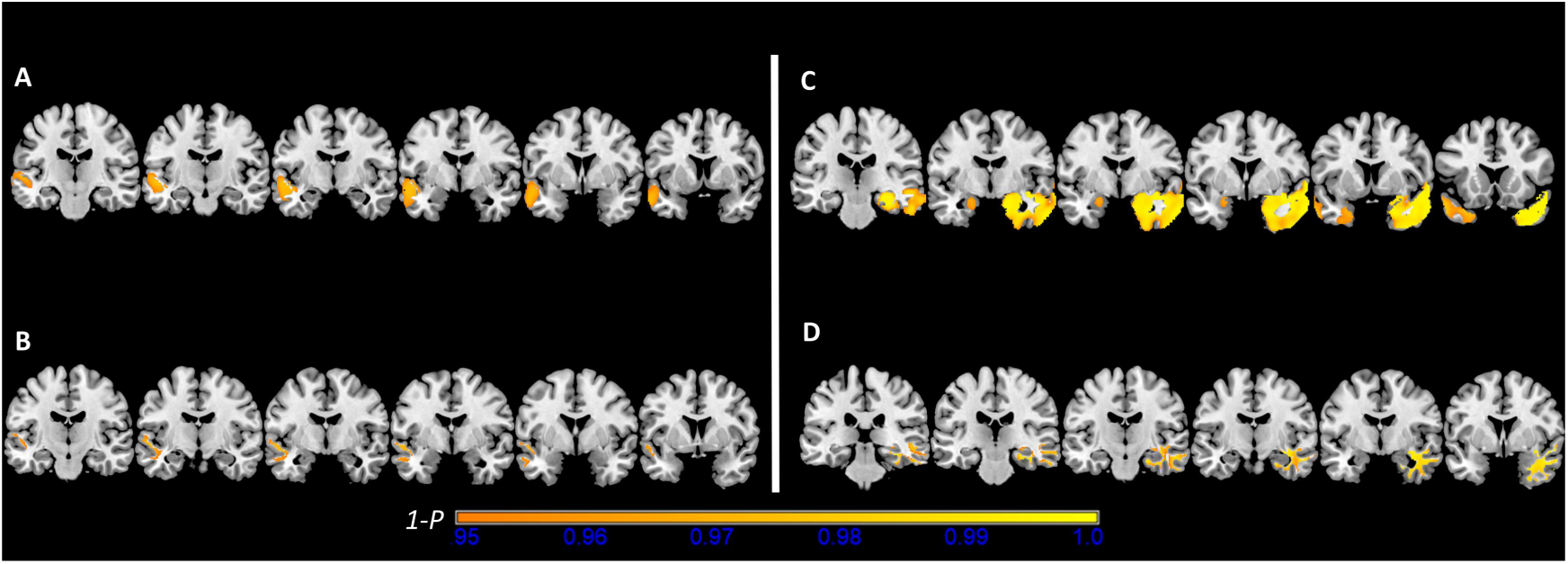
APOE polymorphism modulated atrophy spatial pattern in temporal lobe after mTBI. (A) mTBI APOE ε4 carriers showed severe atrophy at the left superior temporal gyrus and middle temporal gyrus. (B) mTBI APOE ε4 carriers showed severer atrophy of white matter underlying the left superior temporal gyrus and middle temporal gyrus. (C) mTBI APOE ε4 non-carriers showed severe atrophy at the bilateral medial temporal areas of grey matter, and extensive areas in the right lateral temporal cortex. (D) mTBI APOE ε4 non-carriers showed severe atrophy at the right temporal white matter. Results are thresholded at voxelwise *P* < 0.05, with threshold-free cluster enhancement (TFCE) correction.

### Longitudinal volume changes and neuropsychological performance

For mild patients carrying APOE ε4 allele, the performance change-ratio of language fluency could be predicted by the atrophy of the superior temporal gyrus and middle temporal gyrus in the gray matter (model 1: *β* = 0.570, *P* < 0.05) rather than in the white matter (*P* > 0.05). We further controlled sex, age and education in this regression model in a step-by-step way. The atrophy of the same regions can still predict the performance change-ratio of language fluency after controlled sex and age (model 2: *β* = 0.560, *P* < 0.05; model 3: *β* = 0.560, *P* = 0.059), but became failure after controlling the education level (model 4: *β* = 0.420, *P* = 0.19). These relations cannot be replicated in the mTBI patients without carrying APOE ε4 in the neither GM nor WM (*P* > 0.05).

## Discussion

In this study, we found that APOE polymorphism modulated the longitudinally spatial atrophy pattern of the temporal lobe in patients with mTBI. The lower JD value across the 6-12 month follow-up period mainly exhibited in the left temporal lobe in mTBI patients than that of HCs. For the patients carrying APOE ε4 allele, there was the longitudinal atrophy in the left superior temporal gyrus and middle temporal gyrus regions, in which the atrophy of the gray matter was capable of predicting predict the performances changes of language function. While, patients without APOE ε4 allele presented the longitudinal atrophy mainly in the bilateral medial temporal lobe and right lateral temporal lobe.

Mild TBI caused the longitudinal brain atrophy in both gray matter and white matter of the temporal lobe during one-year follow-up. The temporal lobe is one of the most insulted areas impacted by TBI[35-37], in which the atrophy can be generally detected after several years following mTBI[38-40]. In the current study, we adopted a deformation-based morphometry method to directly model the rate of brain atrophy in a voxel-wise way. This analysis captures the repeated-measures characteristic in a longitudinal investigation and providing greater sensitivity and reducing biases from ‘asymmetric’ imaging analyses. By using this method, we can detect the significantly progressive atrophy across the 6-12 months post-injury following mTBI.

For patients with APOE ε4 allele, salient atrophy was mainly exhibited in the left superior temporal gyrus and middle temporal gyrus. These areas are reported to the initially salient atrophy areas in the logopenic variant of the primary progressive aphasia (lvPPA) AD phenotype[16] and served as a neuroimaging signature to identify the lvPPA AD onset risk.

APOE ε4 allele carriers are reported to take accounts of nearly 70% in the lvPPA AD phenotype population [41], and is considered as an important genetic determinant for the onset of the lvPPA AD. Previous study reported that TBI patients carrying APOE ε4 allele showed several neurobehavioural disturbances which was inferred correlated with the structural deterioration in the temporal lobe[42]. The current findings provided further evidence that the longitudinal atrophy of the left superior temporal gyrus and middle temporal gyrus could predict the language dysfunction in mTBI APOE ε4 carriers. Additionally, one recent study reported that the AD genetic factor interacting with mTBI caused the decreased cortical thickness in the AD-vulnerable regions[43]. Our findings support that mild TBI is associated with worse neurodegeneration in the lvPPA AD sensitive regions and reduced language deficits in individuals with the genetic risk for Alzheimer’s disease (i.e., APOE ε4 positive).

The extensive atrophy in the MTL area was particularly observed in APOE ε4 non-carriers following mTBI. TBI usually caused structural and functional integrity decrease of the MTL, especially in the hippocampus and related structures[44-46]. Even a severe tau pathology in the MTL could be detected after the mild injury[47]. These neuroimaging evidences suggest that the deterioration in MTL areas is vulnerable to TBI insult. In the present study, we observed the longitudinal atrophy in the MTL regions in mTBI individuals with low risk of AD genotypes. This current result supports previous finding that the development atrophy in MTL regions following TBI is independent of APOE ε4 allele [48] and may has little reference to AD progression post-injury. Instead, such progressive atrophy observed in mTBI APOE ε4 non-carriers possibly indicates the injury related longitudinal deterioration.

There were several limitations in the present study. First, the APOE ε4 allele is termed as the genetic risk factor of AD, which is also exerts a dosage-dependent effect on the risk of AD[49]. Nevertheless, the APOE ε4/ε4 homozygotes only take nearly 10% of AD affected individuals[50, 51]. Our current study was limited to the relative sample of APOEε4 positive mTBI patients, and further study should include larger sample to investigate the dosage effects of APOE ε4 allele on the longitudinal atrophy of the temporal lobe. Another possible limitation is that the APOE ε4 carriers following mTBI showed the longitudinal atrophy in the left superior temporal gyrus and middle temporal gyrus within the initial year post-injury.

This atrophy pattern is characteristic of the initial neurodegeneration pathway for the lvPPA AD. However, such finding should be verified in a long-term observation on the atrophy trajectory after mTBI, which may provide more evidences on the risk of lvPPA AD following mTBI.

In sum, APOE alleles can modulate the progressive atrophy in the temporal lobe following mTBI, especially in the neurodegeneration pathway of the lvPPA AD. These findings underscored that TBI interacted with high genetic risk to induce long-term neurodegenerative consequence of given phenotype of AD.

